# From Laboratory to Industry: Awareness of the translational potential of Extracellular Vesicles. Results of first survey performed by the ISEV Translation, Regulation and Advocacy Committee (ISEV-TRA)

**DOI:** 10.1101/2025.10.31.685739

**Authors:** Uxia Gurriaran-Rodriguez, Benedetta Bussolati, Mario Gimona, Konstantin Glebov, Yu Fujita, Antonio Marcilla, Christian Neri, Qing-Ling Fu, Saumya Das, Stefano Pluchino, Natasa Zarovni, Carlos Salomon, Kenneth Witwer, Juan Manuel Falcon-Perez

## Abstract

Extracellular vesicles (EVs) are critical mediators of cellular communication, with significant potential for disease diagnosis, therapeutics, and other industrial applications. However, translating EV-based innovations into real-world products faces substantial challenges, particularly concerning regulatory frameworks and scientific gaps. To address these issues, the **International Society for Extracellular Vesicles (ISEV)** established the **Translation, Regulation, and Advocacy Committee (ISEV-TRA)** to create an environment that facilitates the translation of EV research into practical societal applications. This reporet presents the results of the first survey conducted by the ISEV-TRA, assessing the degree to which EVs have been translated into society applications, awareness of EV potential, and the current use of EVs across various industry sectors. Respondents included individuals from Academia, Industry, and dual affiliation (Academia-Industry), provided an initial perspective on EV research and its translational status. The survey highlights the growing interest in EV-based products, suggesting that there is an important wave of efforts to develop these products. Several spin-outs created by academic researchers are focusing on EV-based products, a key trend in the field, while large companies are showing increasing interest in EV-based technologies. The expectations and suggestions for the newly established ISEV-TRA underscore the need for a unified approach to accelerate the translation of EV-based technologies into practical applications. The ISEV-TRA is well-positioned to play a key role by providing essential resources, organizing workshops, and promoting interdisciplinary collaboration, ultimately driving the commercialization of EV-based innovations.

## INTRODUCTION

In recent decades, extracellular vesicles (EVs) have emerged as essential mediators of intracellular communication in both physiological and pathological conditions. Acting as natural shuttles, they transport bioactive cargo—such as proteins, microRNAs, and lipids—to distant targets, influencing various biological processes. This intrinsic communication capability, and the cargo composition traceable to original cell and tissue homeostasis alterations, has endowed EVs with significant translational potential in biomedical sciences, particularly in diagnostics, drug delivery, and overall, clinically applicable therapeutics. Numerous patents and clinical trials attest to their promise^1^.

Beyond biomedical applications, EVs are now being explored in novel fields, including cosmetics, food science, dietary supplements, agriculture, and even ecological sustainability. However, despite this growing interest, significant challenges hinder the full-scale translation of these discoveries into industry and society. A major barrier is the reluctance of the industrial sector to invest in EV-based technologies due to unresolved regulatory and scientific gaps. Notably, neither the U.S. Food and Drug Administration (FDA)^2^ nor the European Medicines Agency (EMA)^3^ has approved any EV- or exosome-derived product for clinical use, citing critical knowledge gaps that must be addressed before regulatory approval can be granted. Furthermore, the International Society for Extracellular Vesicles (ISEV)^4^ has released a safety notice stating that all EV- or exosome-based therapies are experimental at this stage and should be used solely for research purposes. Any potential new therapy must be duly approved by an appropriate healthcare regulatory agency, such as the FDA in the United States or the EMA in the European Union.

It is important to note that this publication is not focused on the therapeutic translational aspect, as there is already covered by the Regulatory Affairs and Clinical Use of EV-based Therapeutics Task Force established by the ISEV for that specific purpose. Additionally, the current state of translational EV-based therapies has been extensively reviewed in previous publications^5–10^.

To bridge the gap in EV translational research beyond clinical applications and to promote responsible, regulated innovation and adoption across various sectors, the **ISEV Translation, Regulation, and Advocacy Committee** (**ISEV-TRA**) was established in 2024. The medium-long goal of the **ISEV-TRA** is to collect and identify key gaps and engage stakeholders across the entire value chain, laying the foundation for a comprehensive innovation management, validation and regulatory framework, and advocacy strategy. By doing so, it should facilitate the successful translation of EV-based innovations into final products, regardless of the Industry sector.

Here, we present the results of the first worldwide survey conducted by the **ISEV-TRA** as an initial point to set the pillars for future directions. By collecting and openly sharing these insights, our goal is to assess the current state of EV translation, raise awareness on the evolution of the translational and innovation landscape for EV research and to deep dive into key aspects of the whole innovation value chain, in view of fostering sectoral engagement and academic-industry partnerships to accelerate the safe and effective application of EV-based technologies to health and disease. Additionally, the survey seeks to understand the expectations for this newly formed committee.

### THE SURVEY AND THE SAMPLE TYPE/SIZE

This survey, consisting of 16 questions (both open-ended and multiple-choice), was developed by the **ISEV-TRA**. The full set of answers is provided in **Supplementary Table S1**, while the original survey format is shown in **Supplementary Table S2** and accessible into the ISEV webpage. Since the **ISEV-TRA** was recently established, no prior data exist for retrospective trend analysis.

To maximize participation, ISEV promoted the survey through multiple electronic announcements, including emails to the ISEV mailing list, posts on social media platforms (e.g., Facebook and Twitter), and outreach at the ISEV2024 Annual Meeting.

A total of 146 full or partial responses were recorded. Among respondents, 86% were current ISEV members, 10% were former members, and 4% had never been affiliated with ISEV (**Figure 1A**). Representation across sectors included 121 participants from Academia, 14 from Industry, 9 with dual affiliation (Academia-Industry), and 2 classified as N/A (**Figure 1B**). Of these, 144 responses were complete (i.e., all questions answered). However, two submissions were excluded from the final analysis due to a high number of unanswered questions—one respondent was retired, and another was the owner of an aesthetic clinic— together representing only 1% of the sample.

**Figure 1.**
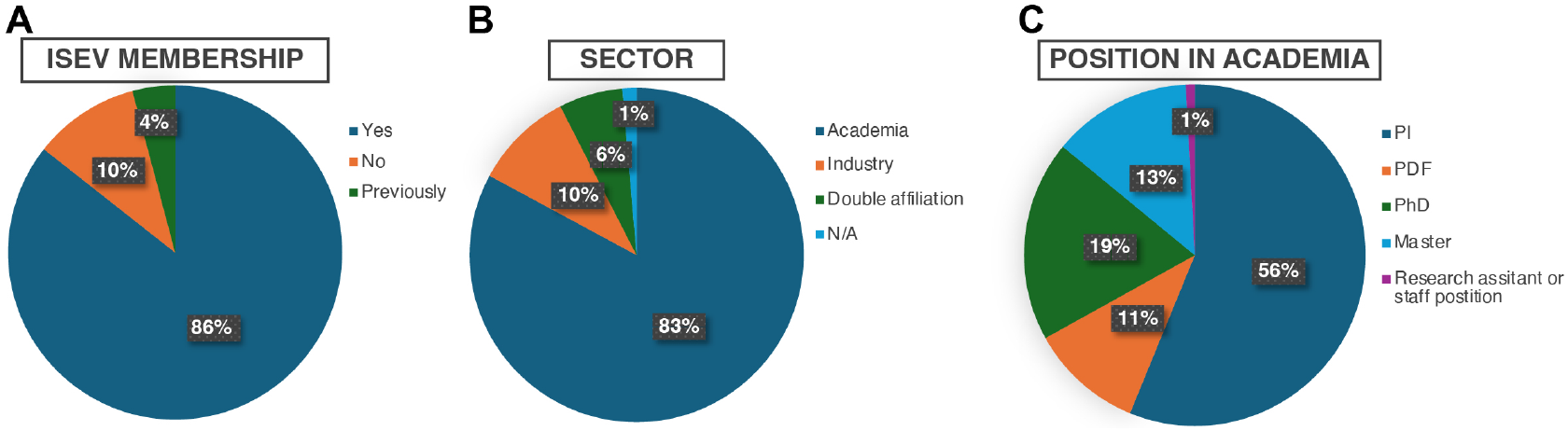
Sample characteristics by type and sectorial distribution. **(A)** Distribution of individuals based on ISEV affiliation. **(B)** Distribution across professional sectors. “Double affiliation” refers to individuals working simultaneously in academia and industry. **(C)** Professional categorization within academia. Data are presented as percentages, based on a total of 146 individuals. All reported percentages are calculated from the respective sample size for each question.

For subsequent analyses, responses were stratified into three categories: Academia, Industry, and Double Affiliation (Academia-Industry). Academia accounted for 83% of respondents, with a balanced distribution across different academic positions (**Figure 1C**). The most represented role was Principal Investigator (56%). In contrast, Industry and Double Affiliation participants were not further stratified by position due to the diversity of terminology for leadership and executive roles, which were the most frequently reported.

This distribution highlights engagement at the senior and executive levels within the surveyed population. The number of respondents per question is provided in the corresponding figure legends, and all reported percentages are based on the respective sample size for each question.

### GEOGRAPHICAL DISTRIBUTION AND SECTOR REPRESENTATION IN EV TRANSLATIONAL RESEARCH

We analyzed the 144 survey responses based on ISEV chapter representation, finding that all three chapters were included. The Europe-Africa chapter had the highest representation (46%), followed by the Americas (33%) and Asia-Pacific (21%) (**Figure 2A**).

**Figure 2.**
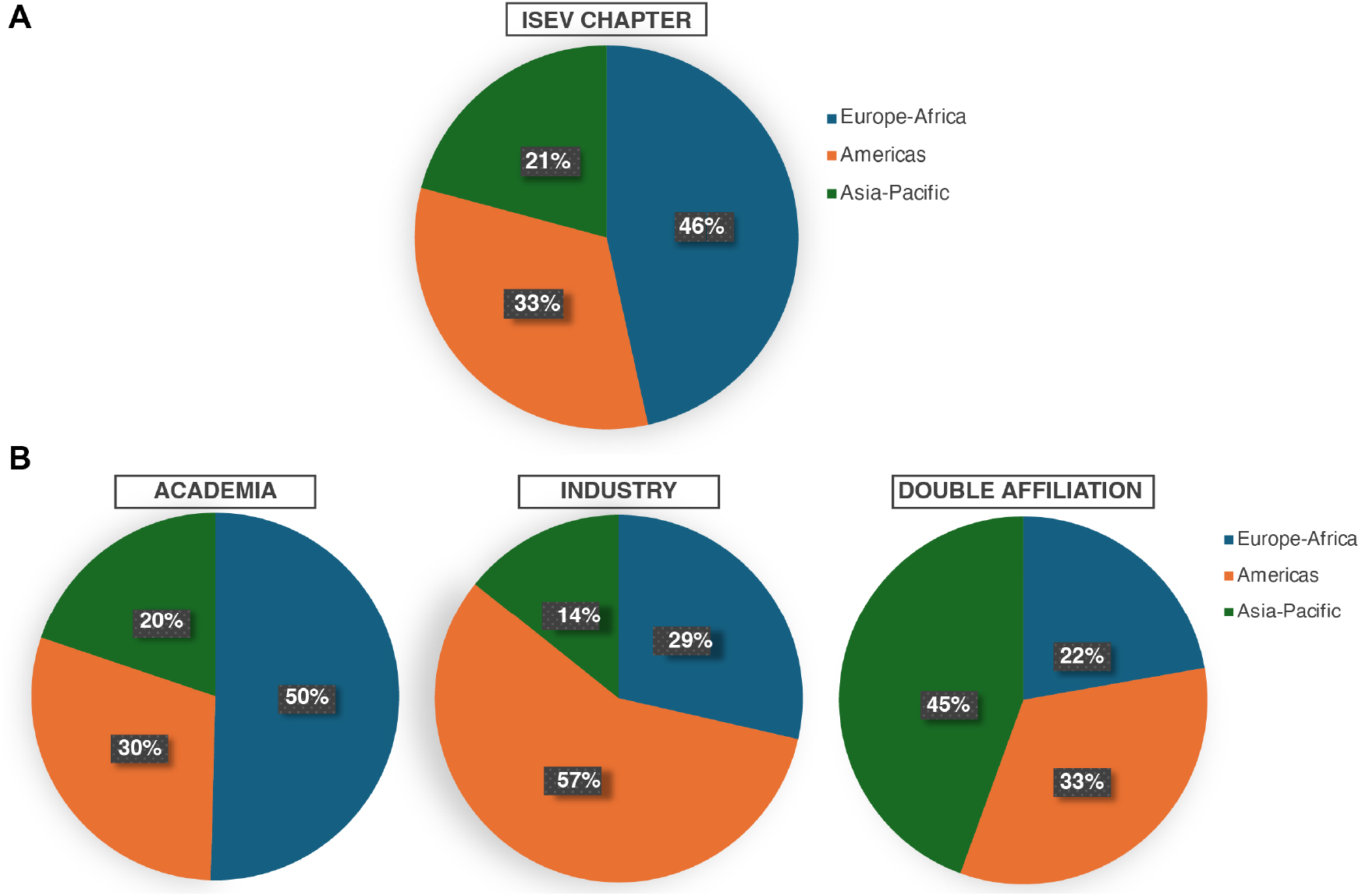
Geographical and sectoral distribution in EV translational research. **(A)** Distribution of individuals across the three ISEV chapters (Europe-Africa, Americas, and Asia-Pacific). **(B)** Sector-specific distribution within ISEV chapters, categorized into academia, industry, and double affiliation. “Double affiliation” refers to individuals working simultaneously in academia and industry. Data are presented as percentages, based on a total of 144 individuals. All reported percentages are calculated from the respective sample size for each question.

Nonetheless, when stratifying the data by sector and geographic region, distinct patterns emerged (**Figure 2B**). Academia was most represented in the Europe-Africa chapter, whereas Industry had a stronger presence in the Americas. Interestingly, respondents with dual affiliation (Academia-Industry) were predominantly from the Asia-Pacific region.

These findings suggest that geographical factors may influence the development and adoption of EV translational research. Despite having the largest share of survey respondents, Europe-Africa appears to have lower engagement in EV translation compared to other regions within the ISEV community. This disparity could be due to many factors including potential regional differences in investment, regulatory environments, and Industry-Academia collaboration for EV-based innovations as it has been already stated by Wang et al^5^. Alternative, these observations could suggest that EV translational research is a trend that is evenly represented across continents where slight differences in percentages may represent differences in sectorial landscapes and/or maturity of EV translational research programs across countries rather than true differences in the level of engagement into innovation and industry. However, given the disproportion of survey respondents from different geographies, such a preliminary conclusion can be skewed by a limited sample size and consequently the suggested trend of this first questionary will need to be further followed by future surveys.

### STATE OF THE ART OF TRANSLATIONAL EV RESEARCH

#### Academic Spin-outs

In this study, academic spin-outs refer to companies founded by academic institutions to develop and commercialize EV-based technologies. Among the 12 spin-outs identified in our sample, 50% originated in Europe, 25% in the Americas, and 25% in Asia-Pacific (**Figure 3A**). Notably, while Industry and Double Affiliation were more prevalent in the Americas and Asia-Pacific, respectively, the Europe-Africa chapter had the highest proportion of spin-outs.

**Figure 3.**
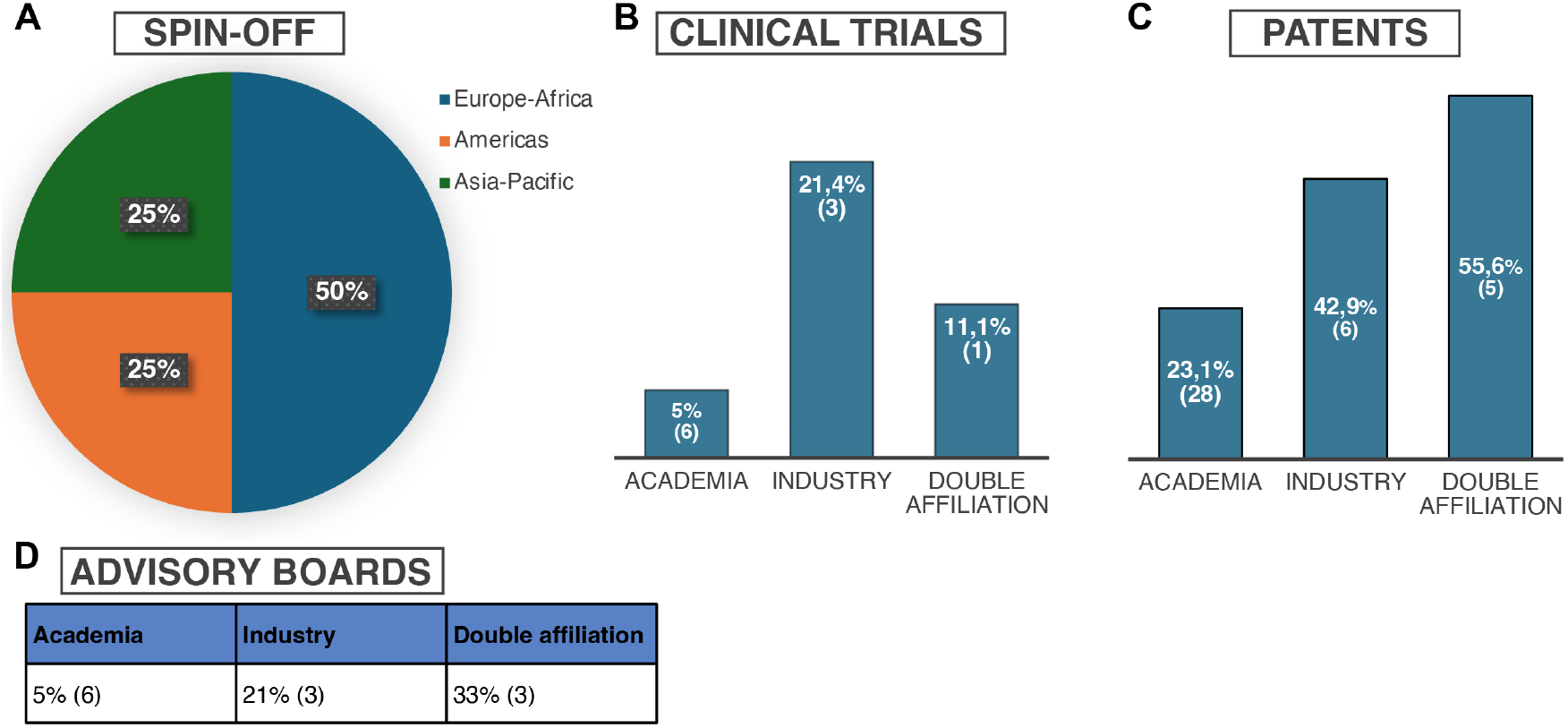
Quantification of EV translational research activities. **(A)** Distribution of spin-offs across the three ISEV chapters (Europe-Africa, Americas, and Asia-Pacific), presented as percentages. **(B)** Number of clinical trials (related to or utilizing EVs) by professional sector (academia, industry, and double affiliation), presented as percentages with absolute numbers in brackets. **(C)** Number of patents by professional sector, presented as percentages with absolute numbers in brackets. **(D)** Advisory board affiliations by professional sector. Data are presented as percentages with absolute numbers in brackets, based on a total of 144 individuals. All reported percentages are calculated from the respective sample size for each professional sector.

Spin-outs represent the early stages of translational research and may also reflect the differences in the availability and enthusiasm of private funding entities such as venture funds to invest in early-stage new companies in the Americas and Asia-Pacific compared to Europe and Africa. Although this is a first survey, the results suggest a favorable environment for academic EV translational research in Europe. However, while Europe and Africa lead in the number of early-stage spin-outs, they lag in established EV-translational industries. This discrepancy highlights the need for increased industry engagement, investment, and supportive mechanisms to successfully translate EV-based innovations from Academia to commercially viable enterprises.

#### Advisory Board Participation

Participation in advisory boards, which typically involves providing expert guidance on scientific, regulatory, and business strategy, was low across all sectors. Only 12 respondents reported serving on an advisory board, with 3 (21%) from Industry, 3 (33%) with hybrid (Academia-Industry) affiliations, and 6 (50%) from Academia (**FIGURE 3D**). Given the low absolute numbers, further stratification by ISEV chapter was not performed.

#### Clinical Trials and Patents

Clinical trials involving EVs remain in early stages, with only a few trials that have reached phase II-III. The reason for such an early stage in clinical validation can be a quality manufacturing bottleneck still staggering to ensure larger trial batches, besides uncertainties in mechanisms of action and competitive advantages, including the cost-effectiveness, with respect to state-of-art options.

According to our survey results, only 7.6% (10 out of 144) of respondents reported involvement in EV-related clinical trials, with 6 from Academia (5%), 3 from Industry (21.4%), and 1 with a Double Affiliation (11.1%) (**Figure 3B**). Notably, the Industry sector shows the highest proportional involvement. Double Affiliation is second, suggesting that for academic researchers to transfer their data to newly created start-ups (e.g. as a founder or else) is a noticeable trend. However, the overall low percentage across all sectors suggests that, as expected, EV-based products are far from industrial commercialization.

Despite the limited clinical validation, the high number of patents across all sectors indicates the pushing in the initial steps of the EV translation (**Figure 3C**). This supports the need for guidelines in understanding and navigating the legal and knowledge protection and transfer frameworks to inform and promote the development and commercialization of EV-based technologies.

### CURRENT TRENDS AND FUTURE DIRECTIONS IN EV RESEARCH

Our analysis stratified EV research by sector, revealing distinct trends in research focus and future interest (**Figure 4**). Fundamental EV-biology research remains predominant within Academia, reinforcing its role as the primary driver for advances in EV space. In contrast, EV research related to Diagnosis and Therapy is evenly distributed across Academia, Industry, and Double Affiliation, indicating a shared interest in clinical applications.

**Figure 4.**
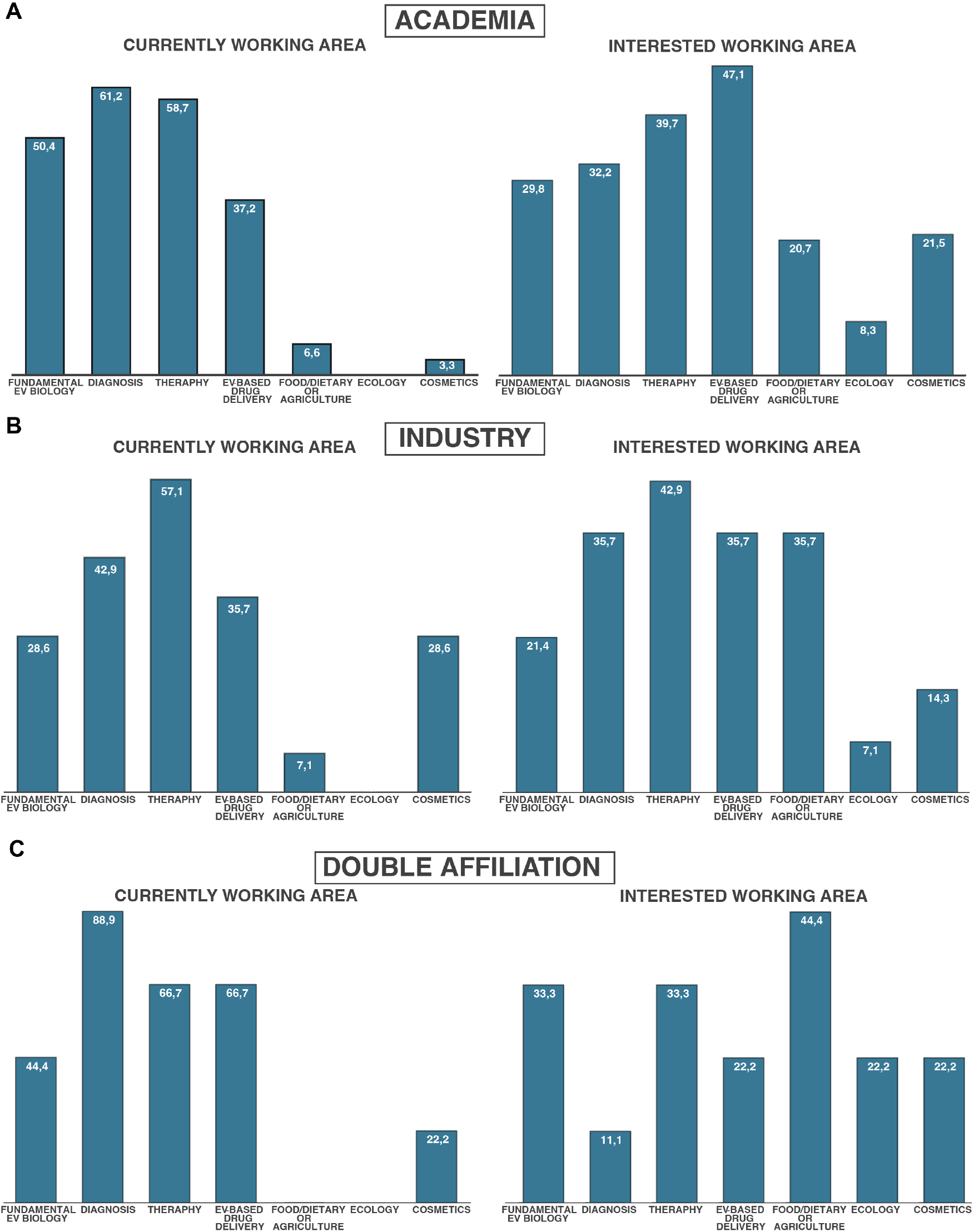
Current trends and future directions in EV research, stratified by professional sector. **(A)** Academia. **(B)** Industry. **(C)** Double affiliation, referring to individuals working simultaneously in academia and industry. Data are presented as percentages, based on a total of 144 individuals. All reported percentages are calculated from the number of positive responses for each expertise field within each professional sector.

Notably, Food/Dietary and Agricultural applications of EV research are currently minimal across all sectors from Academia to Industry, yet there is a strong interest in exploring these areas further. A similar trend is observed for environmental applications, which are absent in our dataset but recognized as an area of potential development. These findings suggest that while EV applications beyond Medicine are overlooked, researchers and Industry stakeholders are considering broad them.

Interestingly, the cosmetic field shows a divergent pattern: while Academic researchers demonstrate a high level of interest, this enthusiasm is apparently not mirrored in the Industry or Double Affiliation respondents. This finding is somewhat surprising as commercial initiatives may benefit from less stringent regulatory hurdles and commercialization challenges in cosmetics. On the other hand, this first survey results, evidence differing and moving research priorities within and between sectors. Often there is a consistent continuity and overlap in research piloting the use of EVs in cosmetics, with their applications in regenerative medicine or as cosmeceuticals. Several clinics offers this kind of “treatment”, but efficacy is not demonstrated, because product content and safety are unclear (e.g., EVs from bacteria).

Overall, our data suggest that while traditional areas like Diagnosis and Therapy maintain a balanced presence across sectors, novel applications such as Food, Agriculture, and Environmental appear as emerging as areas of interest, potentially shaping the future landscape of EV translational research.

### AWARENESS AND ENGAGEMENT IN EV TRANSLATIONAL RESEARCH

The data from the survey highlights significant gaps in awareness and engagement with EV translational research resources among Academia, Industry, and those with double affiliation (**TABLE 1**). While Industry exhibits the highest awareness and participation across most categories (92.86% for both using and selling EVs for research, and 78.57% for public funding awareness), Academia showed a lesser understanding particularly in private funding opportunities such us private investors (only 14.05%). Double-affiliated individuals show moderate awareness levels, often aligning more with Industry than with Academia.

**Table 1.**
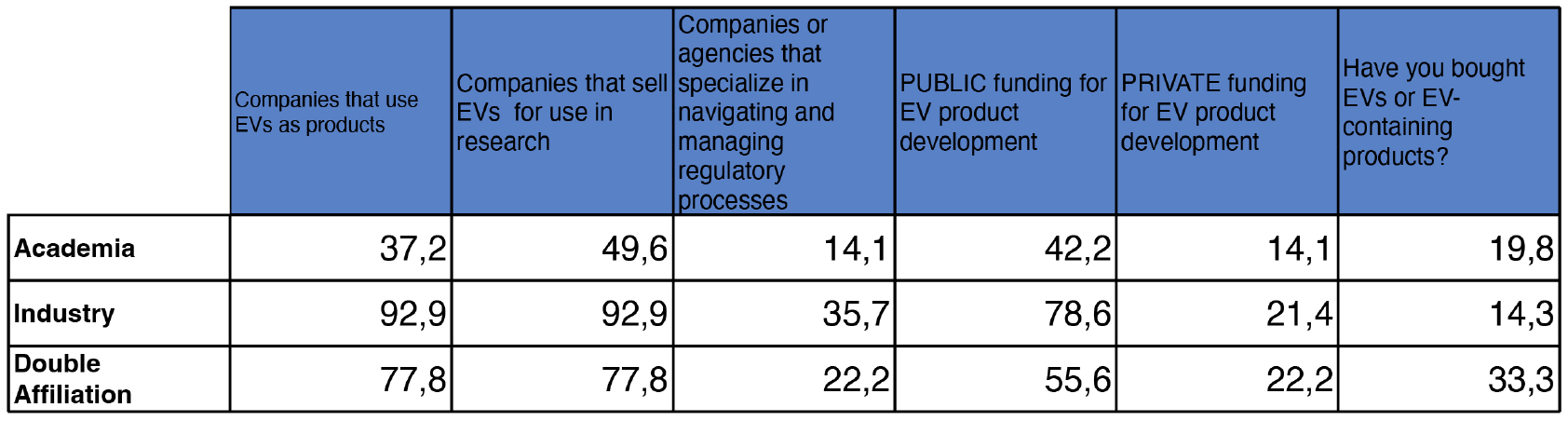
Awareness and engagement in EV translational research, stratified by professional sector. Data are presented as percentages, based on a total of 144 individuals who responded to each question. All reported percentages are calculated from the number of positive responses within each professional sector.

Notably, awareness of regulatory agencies specializing in EV translational research is low across all groups (35.71% in Industry, 14.05% in Academia), underscoring the necessity of initiatives aimed at educating researchers on available resources. Furthermore, the relatively low percentage of respondents who have purchased EV-containing products for research purposes (ranging from 14.29% in Industry to 33.33% in double affiliation) suggests a gap between research and practical applications. These findings suggest the importance of increasing awareness and accessibility of translational research resources to bridge EV research and clinical or commercial application.

### MAJOR OBSTACLES FOR EV TRANSLATION RESEARCH

After EVs are demonstrated to represent key innovative sources of targets, drugs, or biomarkers (e.g., to fight stress and disease), translational research and R&D on EVs still face numerous challenges rooted in the large heterogeneity of EV subtypes (biogenesis pathways, secretion rates, fate) and the technical difficulties associated with isolating specific EV subtypes from other nanoparticles.

One of the main challenges in academia is the lack of a structured Quality Management System (QMS), which compromises the reproducibility and reliability of results. This is particularly critical for regulatory approval, as clinical applications require consistent and reproducible data to ensure patient safety and efficacy. The heterogeneity of EV populations makes their study complex, as different subtypes can exhibit varying biological functions. The lack of standardized isolation and purification methods, along with issues of reproducibility and scalability, results in trade-offs between selectivity and yield. Additionally, a lack of validated reference materials and commercially available antibodies impairs the ability to characterize EVs reliably. Overall, there is a pressing need for robust characterization techniques, quality control measures, and validated biomarkers to ensure the consistency and efficacy of EV-based therapies.

In the Industry sector, scalability and regulatory uncertainty are the dominant issues. Manufacturing EVs at a large scale while maintaining consistency and potency is a significant challenge, particularly under Good Manufacturing Practice (GMP) conditions. The complexity of upstream and downstream processing requires optimization to ensure reproducibility.

Furthermore, EV-based therapies do not fit neatly into existing biologics or gene therapy categories, unlike more established biological products such as cell-free DNA-based diagnostics, antibody therapies, or gene therapies. Additionally, our understanding of EV pharmacokinetics, biodistribution, long-term stability, and immunogenicity remains limited, thereby hindering progress in both therapeutic and diagnostic applications. Despite the pressing need for standardization in diagnostics, with rigorous assay validation required to ensure clinical reproducibility and compliance with existing GLP or CLSI guidelines, regulatory bodies face difficulties in defining the safety and efficacy criteria for EV-based treatments. As a result, their regulatory pathway remains unclear, affecting investment and confidence in the field. Additionally, the evolving nature of the field means that regulatory guidelines struggle to keep pace with technological advancements, leading to inconsistencies in approval processes across different countries. The design and progression of EV processes, products, and trials will benefit from cross-compliance with more mature regulatory and clinical guidelines for state-of-the-art advanced medicines, tailored to a particular indication of use in which a single EV therapeutic is proposed. Specifically, in the cosmetic field, while some companies market EVs-derived products, the scientific rigor in this space is often lacking, further complicating industry credibility.

Besides the unclear path to regulatory approvals, the EV-based therapeutics often lack the clear path to profitability and adoption. As for every product development, the process should initiate from a clear identification of an unserved clinical or consumer need that is clearly uncovered by state-of-art solutions. The unicity of EV based technologies to provide one is still unclear. The early matching of the proposed EV-derived material to a desired product profile would aid the transfer to investors and industry-supported development. The early thought of reproducibility and economic sustainability must be evermore included if we want the EV translation to progress and thrive

Ultimately, while all three sectors acknowledge the need for improved standardization, reproducibility, and regulatory clarity, the specific challenges vary. Academia focuses on biological understanding and methodological inconsistencies, Industry grapples with scalability and regulatory uncertainty, emphasize the need for validated quality control. Overcoming these barriers will require multidisciplinary collaboration to establish robust guidelines, demonstrate clinical success, and secure sustained investment in the field.

### EXPECTATIONS FROM ISEV TRANSLATION, REGULATION, AND ADVOCACY COMMITTEE

The data of this first survey reveals a demand across all sectors for structured information and resources to support EV translational research (**TABLE 2**). The most universally sought-after resources include a checklist of what is needed to translate an EV product (68.6% in Academia, 64.3% in Industry) and reports on the EV marketplace and degree of product translation, with Academia showing slightly higher interest than the others.

**Table 2.** Expectations from the ISEV Translation, Regulation, and Advocacy Committee. Data are presented as percentages, based on a total of 144 individuals who responded to each item. All reported percentages are calculated from the number of positive responses within each professional sector.

|  | Reports and graphics of the overall EV marketplace | Reports and graphics of the degree of translation of existing EV products | A checklist of what is needed to translate an EV product | List of companies selling material/ instruments of interest for EV research, | List of companies selling material/ instruments of interest for EV clinical applications | List of clinical trials based on EVs | List of patents based on EVs | List of EV products that are already on the market | Accounts of how previous products received regulatory approval | Opinions/ interviews with experts pointing to possible opportunities for EVs in new sectors |
| --- | --- | --- | --- | --- | --- | --- | --- | --- | --- | --- |
| **Academia** | 56,2 | 54,5 | 68,6 | 44,6 | 28,9 | 57,0 | 38,0 | 47,1 | 32,2 | 19,8 |
| **Industry** | 50,0 | 50,0 | 64,3 | 28,6 | 21,4 | 57,1 | 42,9 | 42,9 | 71,4 | 14,3 |
| **Double Affiliation** | 77,8 | 88,9 | 55,6 | 22,2 | 22,2 | 33,3 | 33,3 | 11,1 | 77,8 | 66,7 |

However, distinct sectoral differences emerge in the last three categories. Accounts of previous regulatory approvals are highly valued in Industry (71.4%) and among double-affiliated individuals (77.8%), but Academia showed less interest (32.2%), suggesting that Industry professionals and those with cross-sector experience are more focused on regulatory issues. Similarly, interest in patents and clinical trial data is higher in Industry (42.9% and 57.1% respectively) than in double-affiliated individuals (33.3% and 33.3% respectively), while Academia holds an intermediate position.

Notably, Academia and Industry show limited interest in expert opinions on new EV opportunities (19.8% and 14.3%, respectively), whereas double-affiliated individuals express significantly higher interest (66.7%). This suggests that individuals straddling both research and commercial sectors recognize the value of forward-looking insights more than those confined to a single sector.

Additional comments from the survey, both Academia and Industry, underscore the critical role of ISEV in guiding the translational and regulatory landscape of EV research. Academic researchers emphasize the necessity of regular updates on the committee’s progress, expanding global engagement, and organizing structured regulatory discussions, particularly in alignment with existing nanoparticle guidelines. They also highlight the importance of market reports, clinical production facility listings, and infectious disease applications as essential resources for the field. From the Industry perspective, there is a clear demand for commercialization strategies, regulatory clarity, and workforce training initiatives to ensure a standardized and scalable approach to EV-based products.

Overall, the data highlights the importance of creating, organizing, and sharing comprehensive knowledge resources that address regulatory, market, and translational challenges. By making this information widely accessible, the **ISEV-TRA** committee can bridge existing gaps and facilitate more effective EV product development and commercialization.

### BEYOND CLINICAL TRANSLATION: EXPANDING ISEV’S SCOPE IN EV

The ISEV has long been at the forefront of clinical translational EV research, driving pilots and advancements in EV-based therapeutics and diagnostics. However, the potential of EVs extends far beyond the clinical realm. As the field matures, there is a growing interest in non-clinical applications, such as cosmetics, food and agriculture, and environmental sciences as shown in **Figure 4**. These emerging areas represent promising avenues for innovation, yet they remain underexplored within the EV community.

The data strongly supports the idea that ISEV should engage in discussions on all three application areas: cosmetics, food/agriculture, and environmental applications (**TABLE 3**). Notably, cosmetics emerge as a widely supported topic across all sectors, highlighting a shared interest in exploring EV-based innovations in skincare and beauty products such as anti-ageing and hair growth products. Of note, the regulatory requirements for cosmetics are different across the globe, and are also evolving, becoming more stringent in some geographies such as Europe, especially for the products that are at the intersection between “white” cosmetics, and cosmeceuticals or therapeutics, driving more companies in the space to adopt clinical grade manufacturing processes.

**Table 3.** Interest in emerging EV research fields, including cosmetics, environmental applications and food industry & agriculture, stratified by professional sector. Data are presented as percentages, based on a total of 144 individuals who responded to each item. All reported percentages are calculated from the respective sample size for each professional sector.

Cosmetics
|  | YES | NO | NOT SURE |
| --- | --- | --- | --- |
| Academia | 65,3 | 29,8 | 5 |
| Industry | 71,4 | 21,4 | 7,1 |
| Double Affiliation | 66,7 | 22,2 | 11,1 |

Enviromental applications
|  | YES | NO | NOT SURE |
| --- | --- | --- | --- |
| Academia | 65,3 | 29,8 | 5 |
| Industry | 50,0 | 21,4 | 28,6 |
| Double Affiliation | 66,7 | 33,3 | 0 |

Food industry and agriculture
|  | YES | NO | NOT SURE |
| --- | --- | --- | --- |
| Academia | 71,1 | 25,6 | 3,3 |
| Industry | 85,7 | 0 | 14,3 |
| Double Affiliation | 66,7 | 33,3 | 0 |

However, Industry shows a distinct preference for food and agricultural applications, aligning with previous findings on emerging translational research priorities. This suggests that companies which participate in the survey recognize a strong commercial and functional potential, along with possible competitive advantages for EVs in these sectors, that are prompted by the green and sustainability agendas. In contrast, environmental applications receive less support from respondents coming from Industry, indicating a gap in interest or perceived market viability, likely due to the still lacing pilots and for-long neglected opportunity in this sector.

These findings reinforce the idea that EV translational research has significant untapped potential, particularly in the food-dietary sector. Given the Industry’s high level of interest, ISEV could play a crucial role in facilitating discussions, fostering collaborations, and more important ensuring that EV research is effectively aligned with commercial and regulatory opportunities in this emerging field. It is also likely that there are other overlooked opportunities for EVs (i.e veterinary) that are not covered by our first survey but will be mapped and explored in the future.

Understanding how EVs research can be integrated into diverse industries requires a multidisciplinary approach that bridges fundamental research with commercialization pathways. By broadening its focus to include these translational opportunities, ISEV can play a pivotal role in fostering discussions, building up a common language and a ground for an open exchange and collaboration, guiding regulatory considerations, and ensuring that EV research reaches its full potential across various sectors.

## CONCLUSIONS

This survey represents the first effort from the **ISEV-TRA** to analyze and share the state of the art in EV Translational Research, providing valuable insights into the community’s needs and priorities. Overall, these data highlight the willingness from both sectors, Academia and Industry, to collaborate with ISEV reinforcing the necessity of establishing clear translational pathways, setting regulatory benchmarks, and facilitating interdisciplinary knowledge exchange.

Academics largely seek updated guidelines, standardized protocols, and clearer regulatory framework, along with expanded EV research into new areas. In contrast, Industry and dual-affiliated respondents underscore the importance of EVs manufacturing standards and regulatory guidance to bring EV-based products to market. Both groups share an interest in collaborative efforts and ISEV led standardization, but Academia leans toward fundamental research support, while Industry focuses on practical routes for product development and clinical translation.

Importantly, the **ISEV-TRA** aims to address the needs of the community by providing access to updated guidelines, protocols, and regulatory information. More broadly, **ISEV-TRA** aims to serve as a focal point for Academia and Industry to engage in discussions about best practices and regulatory challenges, as expressed by the participants in the survey. The **ISEV-TRA** should not be just a facilitator but a necessary driving force in organizing, validating, and disseminating essential information on regulatory pathways, market trends, and product development strategies. Ultimately, without such organized efforts, the risk remains that promising EV innovations may struggle to reach clinical and commercial implementation. This reinforces the need for this initiative to shape the future of EV translation across biomedical, industrial, and emerging applications.

### PERSPECTIVES

The **ISEV-TRA** is uniquely positioned to bridge the gap between EV research and industry by proactively gathering and disseminating critical information to the research community to navigate the complexities of EV commercialization more efficiently. This includes providing essential resources such as market reports, regulatory guidelines, and commercialization strategies, thereby equipping researchers and industry professionals with up-to-date, practical knowledge. Beyond knowledge sharing, the committee will actively work towards developing essential tools for legislation, regulation, and advocacy, establishing ISEV as the primary resource for EV translation.

To facilitate this translation further, the **ISEV-TRA** aims to gather all this information in a database or website, as well as host workshops at the annual ISEV meeting, bringing together experts from diverse fields—including policymakers, capital investors, CEOs, researchers, and industry leaders—to address key translational challenges and foster collaborative problem-solving. As the field evolves, additional events may be organized to meet emerging needs.

By fostering a centralized networking platform connecting researchers, industry leaders, and stakeholders, the committee will enhance the visibility of EV-based innovations and stimulate investment, ultimately accelerating their translation into impactful real-world applications.

## REFERENCE

1. Search for: Other terms: exosome | List Results | ClinicalTrials.gov [Internet]. [cited 2025 Feb 12]. Available from: https://clinicaltrials.gov/search?term=exosome&viewType=Table&page=5

2. Consumer Alert on Regenerative Medicine Products Including Stem Cells and Exosomes | FDA [Internet]. [cited 2025 Feb 12]. Available from: https://www.fda.gov/vaccines-blood-biologics/consumers-biologics/consumer-alert-regenerative-medicine-products-including-stem-cells-and-exosomes

3. Scientific recommendations on classification of advanced therapy medicinal products | European Medicines Agency (EMA) [Internet]. [cited 2025 Feb 12]. Available from: https://www.ema.europa.eu/en/human-regulatory-overview/marketing-authorisation/advanced-therapies-marketing-authorisation/scientific-recommendations-classification-advanced-therapy-medicinal-products

4. Patient information and safety notice: extracellular vesicles/exosomes and unproven therapies [Internet]. [cited 2025 Feb 12]. Available from: https://www.isev.org/patient-information-and-safety-notice--extracellular-vesicles-exosomes-and-unproven-therapies

5. Wang C, Tsai T, Lee C. Regulation of exosomes as biologic medicines: Regulatory challenges faced in exosome development and manufacturing processes. Clin Transl Sci. 2024 Aug 8;17(8).

6. van Niel G, Gazeau F, Wilhelm C, Silva AKA. Technological and translational challenges for extracellular vesicle in therapy and diagnosis. Adv Drug Deliv Rev. 2021 Dec;179:114026.

7. Ghodasara A, Raza A, Wolfram J, Salomon C, Popat A. PERSPECTIVE Clinical Translation of Extracellular Vesicles. 2023; Available from: 10.1002/adhm.202301010

8. Wiklander OPB, Brennan MÁ, Lötvall J, Breakefield XO, EL Andaloussi S. Advances in therapeutic applications of extracellular vesicles. Sci Transl Med. 2019 May 15;11(492).

9. Cheng K, Kalluri R. Guidelines for clinical translation and commercialization of extracellular vesicles and exosomes-based therapeutics. Extracellular Vesicle. 2023 Dec;2:100029.

10. Cheng CA. Before Translating Extracellular Vesicles into Personalized Diagnostics and Therapeutics: What We Could Do. Cite This: Mol Pharmaceutics [Internet]. 2024;21. Available from: 10.1021/acs.molpharmaceut.4c00185

